# Cluster-free annotation of single cells using Earth mover’s distance-based classification

**DOI:** 10.1101/2024.03.18.585613

**Authors:** Rikard Forlin, Pouria Tajvar, Nana Wang, Dimos Dimarogonas, Petter Brodin

## Abstract

Grouping individual cells in clusters and annotating these based on feature expression is a common procedure in single-cell analysis pipelines. Multiple methods have been reported for single-cell mRNA sequencing and cytometry datasets where the vast majority rely on sequential 2-step procedures involving I) cell clustering based on notions of similarity and II) cluster annotation via manual or semi-automated methods. However, as arbitrary borders are drawn between more or less similar groups of cells, one cannot guarantee that all cells within a cluster are of the same type. Further, dimensionality reduction has been shown to cause considerable distortion in high-dimensional datasets and is prone to variable annotations of the same cell when relative changes occur in data composition. Another limitation of existing methods is that simultaneous analyses of large sets of cells are computationally expensive and difficult to scale for growing datasets or metanalyses across multiple datasets. Here we present an alternative method based on calculation of Earth Mover’s Distance and a Bayesian classifier coupled to Random Forest, which annotates one cell at a time removing the need for prior clustering and resulting in improved accuracy, better scaling with increasing cell numbers and less computational resources needed.

## Introduction

Single-cell mRNA sequencing (scRNA-seq) and cytometry technologies have made remarkable progress in recent years, becoming more cost-effective and offering increasingly complex, multilayered readouts from individual cells (1,2). This development has surpassed the impressive trajectory of Moore’s law regarding the number of cells measured (3) and opens new avenues for exploring individual cells in great detail. Single-cell transcripts can uncover regulatory mechanisms such as cell-cell dependencies (4–6), reveal cellular differences between health and disease states (7–10), clarify distinct cellular states (11,12) and determine gene regulatory mechanisms across cell populations (13–16).

To derive meaningful insights from transcriptional data, it is essential to accurately identify the cell types present in a dataset. Prior to scRNA-seq, traditional methods for classifying cell types were based on microscopy, and histopathological descriptions of cellular phenotypes (17). In the field of immunology, the use of cell-surface markers has been important for differentiating cell populations and lineages by fluorescence-activated cell sorting (FACS) (18). With the emergence of scRNA-seq, the transcriptional profile of a cell can be used for its classification (17). However, scRNA-seq introduces a greater degree of uncertainty regarding the information obtained from individual cells due to smaller amounts of starting material and greater technical noise (19–21), as well as shallow sampling of the mRNA content of each cell. Furthermore, since the transcriptional activity of a cell is influenced by its cellular context and environment as well as stochastic factors (3), certain cell types can be challenging to distinguish based on sc-mRNA data alone, leading to the development of CITE-seq which offers combined measurements of both proteins and mRNA-molecules (22).

Currently, the most common way to annotate cell types using scRNA-seq data is to first apply principal component analysis (PCA) to reduce dimensionality of the dataset and highlight the genes that exhibit the most variation. This is followed by a clustering algorithm such as Leiden clustering (23), and finally data visualization by Uniform Manifold Approximation and Projection (UMAP) (24). Subsequently, cells clustered together are annotated based on mean or median expression levels of marker transcripts (25).

While this clustering approach is practical, it tends to oversimplify the intricate biological variability and unique characteristics of individual cells. In other words, it relies on all cells within a cluster being of the same cell type, which cannot be guaranteed (26). The process of reducing data from hundreds or thousands of dimensions to two can cause considerable distortion in high-dimensional datasets (27), affecting both local and global relationships and causing a notable percentage of cells to be incorrectly annotated. Furthermore, common clustering approaches do not take into account the probability of a cell belonging to a certain cell type within a cluster, and thus the level of uncertainty that can arise from e.g. noise in data generation is often ignored in downstream analyses.

The clustering of cells has obvious advantages, as manually annotating scRNA-seq data in a cell-by-cell manner is not a realistic task, and cell phenotypes are not absolute but relative features, determined by contrasting the different populations being studied. As cell clustering can be labor-intensive (26), multiple tools have been developed for automatically annotating cell types (28–30). These rely on the unsupervised clustering of cells, followed by manual or semiautomatic annotation according to canonical marker genes. An accurate tool for automated annotation operating on individual cells, that also scales well with increased data size is therefore required to avoid the limitations of clustering-based annotations. Here, we introduce the BayeLeafClassifier (BLC), an automatic classifier of single-cell data on a cell-by-cell basis offering superior accuracy, efficiency and speed compared to currently available procedures for clustering and annotation.

## Results

### BayeLeafClassifier (BLC) a cell-by-cell annotation strategy

BLC uses Earth Mover’s Distance (31) (EMD) for unbiased feature selection and identification of those most informative. This approach has previously been used to successfully extract differentially expressed transcripts from single-cell transcriptomic data (32–34). The features are then fed into a machine learning-based prediction method for cell annotation, using a naïve Bayesian classifier coupled with a Random Forest classifier. This framework allows BLC to take stochastic fluctuations into account and consider the probability of a cell type expressing a specific transcript at a certain level. This will lead to a measure of the overall likelihood of the cell type assignment for all cells, a feature often ignored in downstream analyses (26). BLC is constructed as a framework for finding the genes most informative for cell classification, where classification can describe cell types, subtypes or states. One key advantage of BLC is its agnostic attitude towards the dataset, eliminating the need for prior knowledge of marker genes when training which simplifies the process of generating classifiers for different purposes, thus enhancing its versatility and ease of use in diverse research contexts. Here, we use BLC to classify fundamental cell types in peripheral blood mononuclear cells (PBMCs). Firstly, training was carried out using publicly available datasets consisting of approximately 400,000 cells from PBMCs and whole blood, followed by benchmarking towards previously unseen data against the cluster-based automatic cell type annotation tool scType (28), and another cell-by-cell classifier, scPred (35), which uses feature selection and projection in a reduced dimensional space by PCA. BLC showed an improvement in correct predictions, spent less time annotating cells than scType and scPred, was resistant to the introduction of noise and down-sampling of datasets, and showed a higher credibility regarding certainty-estimation of its annotations. Additionally, it required significantly less computational resources for both model-training and cell-classification compared to the benchmarked methods, making it suitable for larger scale problems as well.

Given pre-annotated training data, BLC can find the most informative features and trains to classify the grouping of a cell. The general framework of building and running BLC is as follows (**Figure 1a-e**): quality-controlled data is used as input to BLC, which then uses non-linear quantile scaling and EMD to find marker genes that best separate each pair of cell types. EMD provides a distance metric between gene expression distributions across two cell types, where a higher EMD value indicates greater efficacy for a gene to distinguish between these cell types, i.e. a more informative gene. This quickly discards typical housekeeping genes or genes that are shared between two cell types, such as CD3D in CD8^+^ T cells and CD4^+^ T cells (**Figure 1f, left panel)** and leaves specific marker genes such as CD3D (**Figure 1f, middle panel**) or lysozyme (LYZ) (**Figure 1f, right panel**) between CD8^+^ T cells and classical monocytes (CMs). As an example, if an unclassified cell has high CD3D expression, it is more likely that this high expression level comes from a CD8^+^ T cell than a CM. Therefore, BLC can use CD3D as one of a set of genes to separate CD8^+^ T cells *vs* classical monocytes, but not to separate CD8^+^ T cells and CD4^+^ T cells. Using marker genes with the highest EMD-distance for each pair of cell types, we trained a classifier using BLC’s framework to distinguish different gene expression levels in different cell types. When annotating a new cell, BLC considers the marker gene sets that have been identified as the most informative for separation of two cell types. Each cell then goes through a pairwise comparison with all cell types, where the expression level of each marker gene is analyzed by the Bayesian classifier followed by the Random Forest classification. From this, BLC calculates the likelihood that a given cell is one of the two cell types compared. This process allows BLC to determine the most likely cell type for the new cell, thus assigning a probability to each cell with its annotation. Going through all cell type comparisons, BLC determines a cell type based on the highest likelihood of exhibiting a cell-specific gene expression profile. For a more detailed discussion, please see the Methods section.

**Figure 1.**
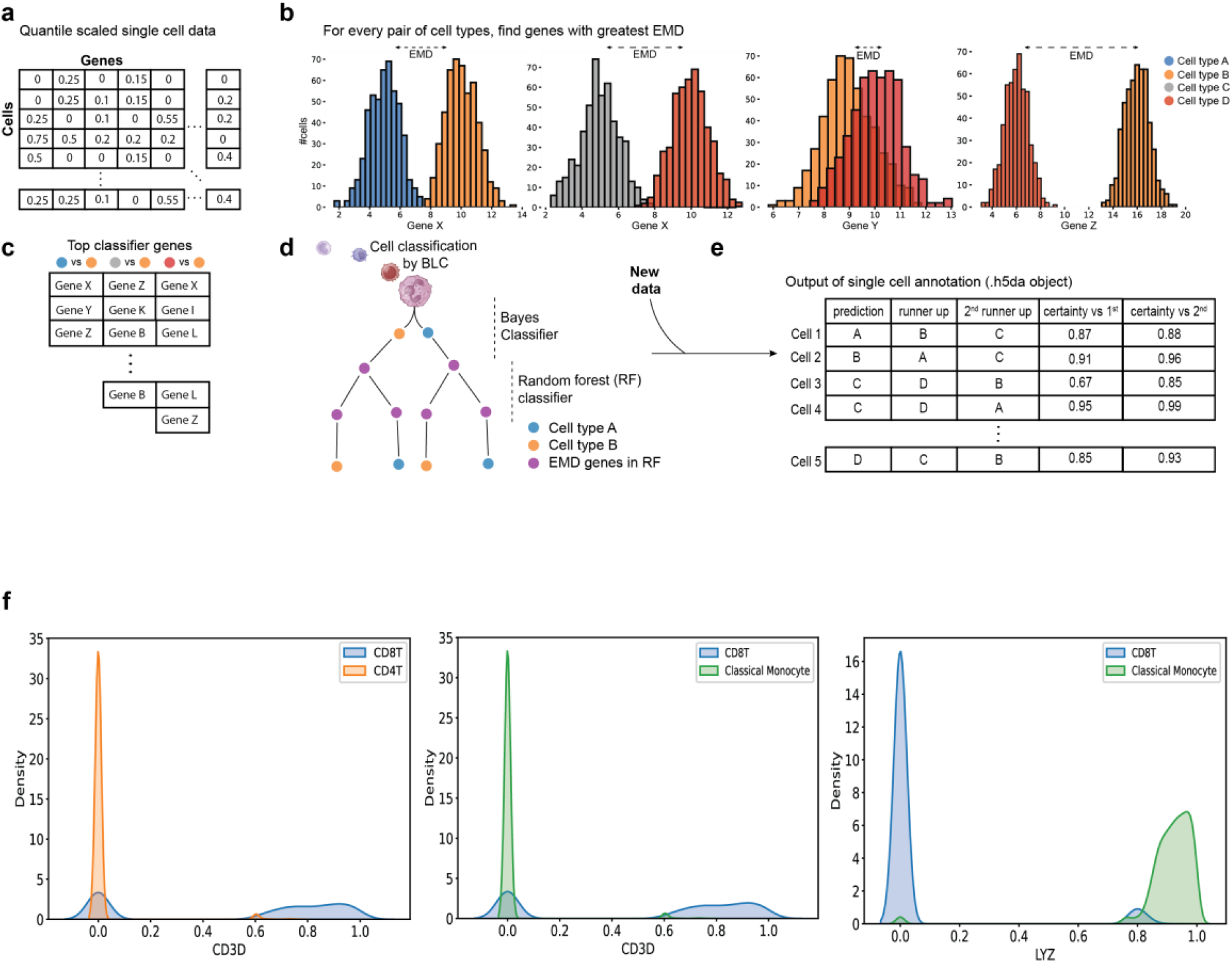
General workflow of BayeLeafClassifier. Quality Controlled (QCed) scRNA-seq data are fed into BLC-workflow and quantile scaled **(a)**. EMD is then performed to find the most informative genes, i.e. genes with the highest distance in their distributions between two cell types **(b)**. These genes are then extracted and fed into the training of the Bayesian-Random Forest classifier BLC **(c, d)**. When classifying new data, BLC takes in an h5ad-object and provides output as metadata in the same h5ad-object **(e). f)** Example of good EMD-marker genes with different distributions in different cell types. CD3D distribution in CD8+T cells and CD4+T cells (left panel), CD3D distribution between CD8+T cells and classical monocytes (middle) and LYZ between CD8+T cells and classical monocytes (right).

In addition to providing the most likely annotation, and the assigned probability for that annotation, BLC also returns the two most likely runners up. These runners up annotations can provide valuable information such as the option to more closely inspect the cells where the classifier displays a higher degree of uncertainty, thereby enhancing the overall transparency, reliability and usefulness of the classification process.

For this article, we built BLC with a maximum of 70 marker genes per cell type (in total, a maximum of 140 genes when the comparison is made) with a minimum EMD-distance of 0.4, a random forest tree with maximum depth of 40, and 200 estimators. The depth limitation and EMD-threshold allowed an optimal balance between capturing nuanced cellular characteristics and ensuring model robustness, thereby preventing overfitting while maintaining high accuracy when classifying test data.

### Benchmarking against available tools

Using our trained classifier, we benchmarked BLC against scType, an automatic cell type classifier based on clustering and scPred, a cell-by-cell classifier with feature selection and projection in a low dimensional space. We selected the micro F1-score as our primary metric for evaluating classification performance, however, calculated macro F1-scores are also available at **Extended Figure 1**. The F1-score is calculated as the harmonic mean of precision and recall across all classes, with micro-averaging to compensate for the class imbalance in cell types (for example, CD4^+^ T cells are much more common than dendritic cells (DCs)). Barplots of original cell annotations are available at **Extended Figure 2**.

For the first test, we used data generated from SmartSeq3 by Hageman-Jensen et al (36), where samples were from healthy controls without any stimulation. Testing SmartSeq3-data, all classifiers scored similarly high F1-scores of 0.93 or 0.94 (scPred) (**Figure 2a**). However, the cluster-based scType was unable to separate B cells from plasma B cells, nor did it separate DCs and plasmacytoid dendritic cells (pDCs).Instead, it classified all plasma B cells as B cells and all the DCs as pDCs, as they were clustered together, resulting in a notably lower macro F1-score of 0.67. scPred scored a macro F1-score of 0.85 (**Extended Figure 1**), while BLC’s F1-score was similar to scPred at 0.84. For the second test, we used a dataset of healthy controls from Lee et al (37). Both BLC and scPred achieved an F1-score of 0.93 while scType scored 0.86 **(Figure 2b)**. While the results were similar for B cells, CD4^+^ T cells, CD8^+^ T cells, CMs, platelets and pDCs, scType had more trouble separating plasma B cells, natural killer (NK) cells and DCs.

**Figure 2.**
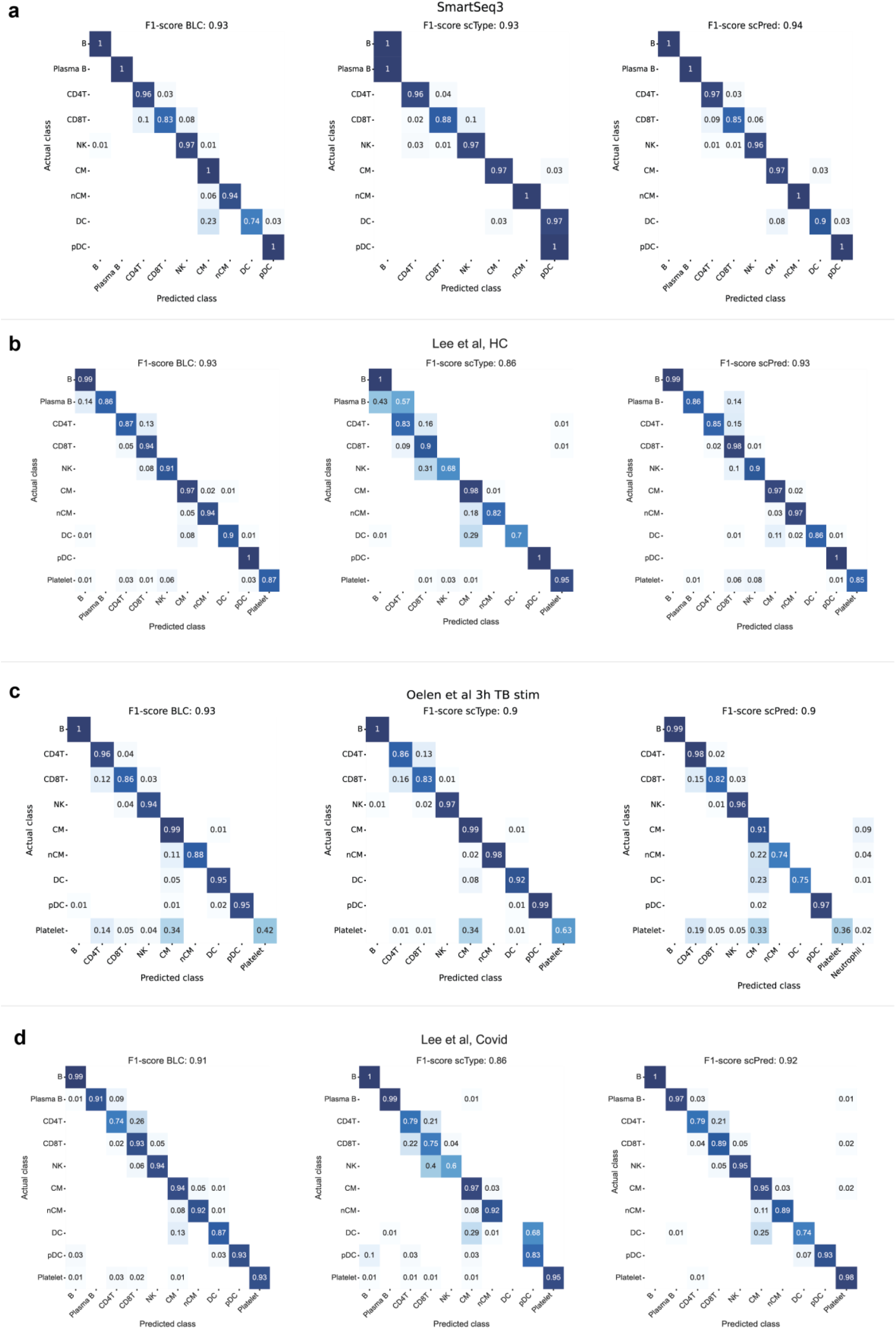
Heatmaps showing micro F1-score and percentage correct annotation of each cell type in dataset: **a)** Smart-seq3 from Hageman-Jensen et al **b)** Lee et al, healthy controls (HC) **c)** Oelen et al, 3 hours TB-stim and **d)** Lee et al, covid patients

We also wanted to challenge the classifiers with activated cells. As a cell transcriptome will change when activated (12), this could impact classification results. Data was obtained from Oelen et al (38), who collected blood samples from healthy individuals and stimulated PBMCs for 3 hours with mycobacterium tuberculosis (MTB). During data generation, two generations of 10X chemistries were applied, both V2 and V3, but only V3 data was classified in the current investigation. BLC achieved an F1-score of 0.93, while both scPred and scType scored 0.9 (**Figure 2c)**. scType had more trouble annotating CD4+T cells *vs* CD8+T cells compared to BLC, however, it did score slightly better regarding the correct annotation of platelets which was difficult for all classifiers (63% for scType, 42% for BLC, 36% for scPred), with all classifiers annotating about a third of them as CMs. scPred had more trouble with CMs, non-classical monocytes (nCMs) and DCs, and labeled ∼2 % of these cells incorrectly as neutrophils.

For a second test with activated cells, we used data from patients with an ongoing COVID-19 infection, also from Lee et al. BLC achieved an F1-score of 0.91, outperforming scType which scored 0.86 **(Figure 2d**), while scPred scored 0.92 being slightly better at classifying platelets and CD4^+^ T cells. BLC performed better regarding the classification of DCs, nCMs and CD8^+^ T cells. scType struggled to accurately annotate NK cells and DCs, the latter being completely missed by scType, while BLC annotated 87% of them correctly.

The fact that scType missed the DC-subset completely for both SmartSeq3 and COVID-19 data, highlights one important issue for clustering approaches, namely the difficulty in separating different yet similar cell types across different cell states, in this case from healthy and COVID-19 affected individuals.

Overall, BLC demonstrates high accuracy and robustness in classifying both perturbed (activated) and unperturbed (inactivated) cells across various datasets. It effectively distinguishes closely related cell populations where traditional clustering methods like scType struggle, highlighting one of the advantages of BLC with increased sensitivity towards less common cell types. Further, BLC demonstrated a higher degree of similarity between the macro F1-scores and the micro F1-scores compared to scType and scPred (**Extended Figure 1**). This suggests that BLC is less biased towards the larger classes and handles the inherent class imbalances found in PBMC data effectively, ensuring that annotations are not disproportionately influenced by the more common cell types in PBMC data.

BLC achieves this high accuracy using ∼140 marker genes to separate different cell types, a significantly smaller number compared to the 2000-3000 marker genes typically used in clustering algorithms to define transcriptional signatures. It does so by effectively using EMD to identify those genes that retain sufficient information to accurately determine cell type. This streamlined, yet efficient selection of marker genes underscores the proficiency of BLC in capturing the essential transcriptional characteristics necessary for precise cell type classification, demonstrating its effectiveness even with a more concise genetic dataset.

### Inconsistent annotations when merging datasets

One issue when performing PCA followed by UMAP for cell type annotation is susceptibility towards different outcomes when the relative composition of the dataset changes. We hypothesized that these inherent properties of clustering algorithms would affect cell annotations by scType to a greater extent than those of our cell-by-cell classifier.

To test this idea, we used the Covid-19 and HC-datasets from Lee et al, and compared annotations generated in separate annotations of the two datasets *vs* the merged dataset. Notably, there was a larger difference between scType annotations for these two conditions, as visualized in UMAP-embeddings (**Figure 3a, b**), but not for BLC annotations (**Figure 3c, d**). For scType, the cell types most affected between conditions were NK-cells, nCMs and DCs, while BLC annotations remained stable across all cell populations **(Figure 3e)**. In total, 11% of cells were inconsistently annotated using scType, while only 2% for BLC (**Figure 3f**). The 2% difference is due to minor variations in the quantile transformation normalization. Moreover, the 2% of inconsistent BLC annotations were characterized by a large bias towards lower uncertainty-levels, compared to first runners up (**Extended Figure 3a**), where 95% were consistent.

**Figure 3.**
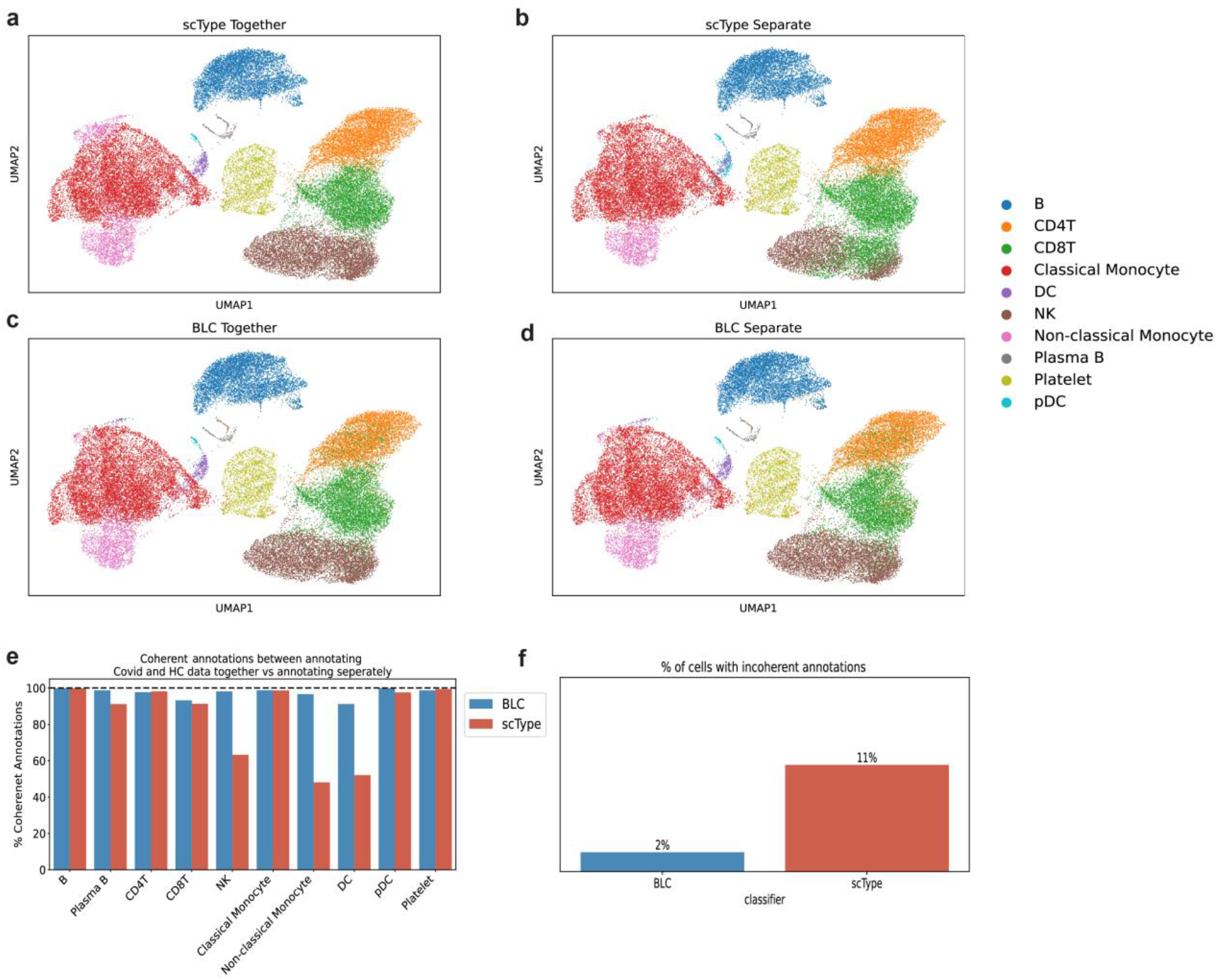
**a-d**: Incoherent annotations of the same cells depending on datasets included. Data from Lee et al from Covid-19 patients and healthy controls (HC) annotated using scType (upper panel) or BLC (lower panel) with both datasets merged (**a** and **c**) or annotated seperately (**b** and **d**). **e**) % of cells per cell type that were coherently annotated in both settings. **f**) Total cells that were incoherently annotated using BLC or scType

To our surprise, scType actually performed better when datasets were merged (**Extended Figure 3b**), as opposed to when they were classified separately (**Figure 2b,d**), and did not become confused by the different states of cells in COVID-patients and healthy controls. However, when classifying new data it is difficult to know beforehand which will give the best result, merged or separate data. In conclusion, BLC gives more consistent results and is a more reliable approach, offering more consistent annotations that are less affected by the relativity of the dataset.

### Inspecting incorrectly annotated cells

To investigate incorrectly annotated cells in more detail, we compared cells that were incorrectly classified by BLC to those incorrectly classified by scType and scPred, to see whether canonical cell marker genes were more aligned with the new or the original annotation, and to better understand the reasons for misclassification among the three different approaches.

Firstly, we compared cells that were wrongly classified and differently annotated between BLC and the other classifiers. The cell type most commonly mislabeled in the COVID-19 dataset from Lee et al were CD4^+^T cells labeled as CD8^+^T cells. As slightly fewer CD4^+^T cells were correctly labeled by BLC, we chose to examine cells labeled CD8^+^T cells by BLC, but where the original annotation was a CD4^+^T cell, and scType/scPred had classified these correctly.

When examining expression of the canonical marker genes that separate CD8^+^ T cells and CD4^+^ T cells (all validated by the Human Protein Atlas (39)), the CD4^+^ T cells that BLC mislabeled as CD8^+^ T cells expressed more of the typical CD8^+^ T cell marker genes (CD8A, CD8B, NKG7, GZMB, GZMA, GZMM, GNLY, KLRD1, CTSW, PRF1) than correctly classified CD4^+^ T cells, as compared to scPred (**Figure 4a, left panel**) and scType (**Figure 4a, right panel**). As another example, we examined DCs that were correctly annotated by BLC, but labeled CMs by scPred. Again, cells mislabelled by BLC were more similar to those marker genes corresponding to BLC annotations than to scPred annotations (**Figure 4b**).

**Figure 4.**
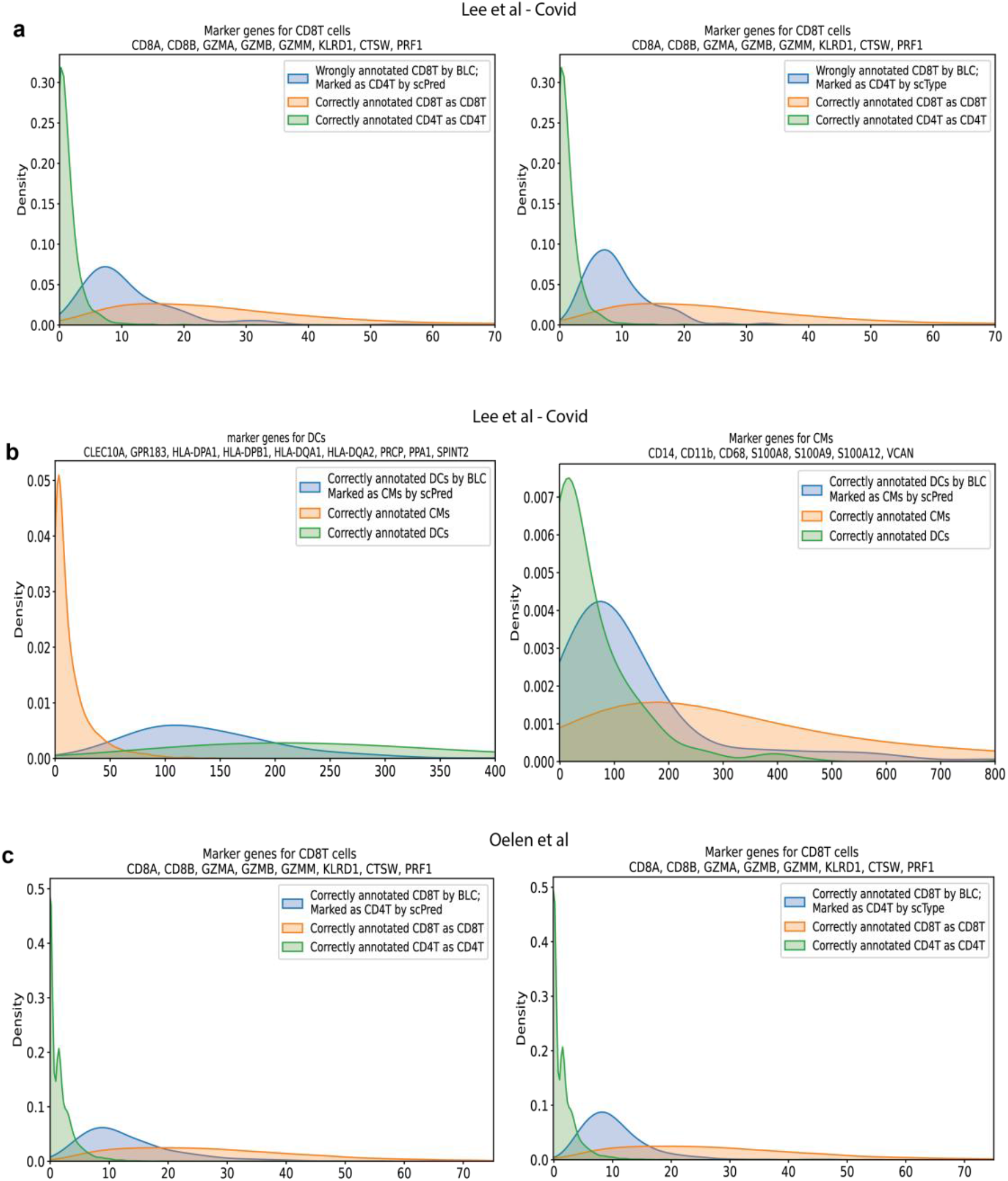
Sums of marker gene distribution in incoherently labeled cells between BLC versus scType and scPred. **a)** CD4+T cells marked as CD8+ T cells by BLC but marked as CD4+ T cells by scPred (left panel) and scType (right panel) in covid data from Lee et al. **b)** Dendritic cells (DCs) correctly annotated by BLC, marked as classical monocytes (CMs) by scPred. Marker genes for DCs (left panel) and CMs (right panel) in covid data from Lee et al. **c)** CD4+ T cells marked as CD8+ T cells by BLC but marked as CD4+ T cells by scPred (left panel) and scType (right panel) in data from Oelen et al.

The above pattern was found across other datasets as well. Looking at cells that BLC correctly classified as CD8^+^ T cells and scPred/scType classified as CD4+T cells, we see that these cells do express more of the typical CD8^+^ T cell marker genes listed above (**Figure 4c**). In total, BLC seem more reliable not only in its correct annotations but also in cases of misclassification, particularly in subsets that are challenging to differentiate, like CD4^+^ T cells versus CD8^+^ T cells or DCs versus CMs.

These findings highlight an interesting feature of our cell-by-cell classifier. Even though BLC was trained on annotations from clustered data (which suffers from some noise due to the nature of the clustering), BLC can classify cells using individual cell characteristics, rather than relying on cluster-based generalizations. BLC can therefore discern subtle patterns in data that traditional clustering approaches might miss. BLC, using EMD to find suitable marker genes, therefore adapts to the inherent heterogeneity of scRNA-seq data to discern individual cellular characteristics.

### Properties of BLC – certainty levels, runners up and time allocation

As BLC also returns a certainty level with its annotations, we noticed that incorrectly labeled cells were generally assigned a lower certainty compared to those correctly annotated. As illustrated in **Figure 5a**, the certainty levels for correctly annotated cells display a pronounced skewing towards higher values, whereas the certainty distribution for incorrect annotations displays a skewing towards lower values. scPred also returns a probability for cell type assignment, but its certainty levels showed a pronounced skewing towards higher values, both when correct and incorrect (**Extended Figure 4a**). This suggests that BLC incorporates a robust self-assessment that scPred is lacking, which is effectively calibrated to reflect a lower confidence in its predictions when approaching the limits of its classification accuracy, thereby providing a reliable indicator of the probability of correctness in its annotations. This characteristic enhances the utility of the model, allowing users to identify and inspect potentially ambiguous cases in greater detail.

**Figure 5.**
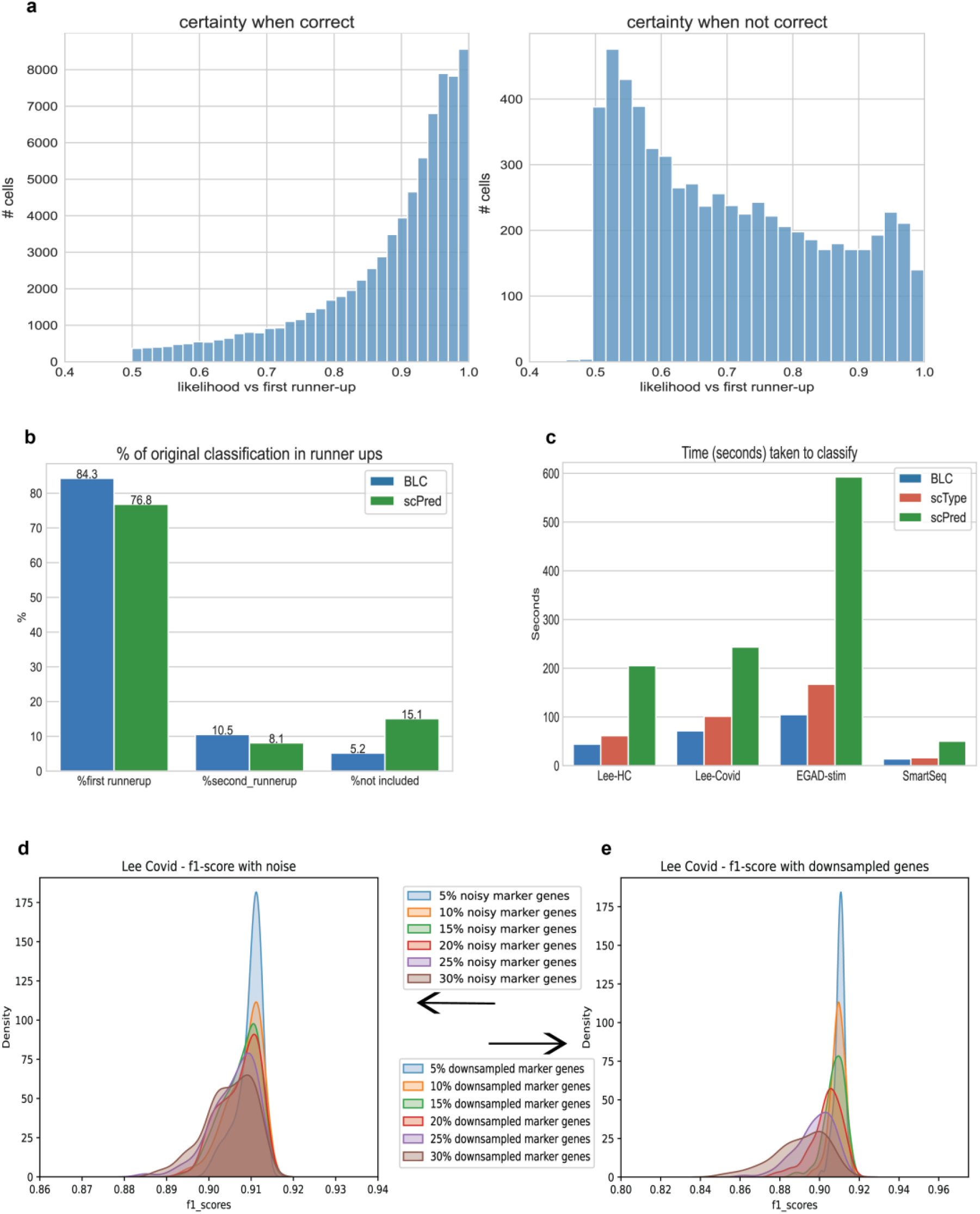
**a)** Certainty levels for BLC when correct (left) and incorrect (right). Likelihood of final label shown in comparison to likelihood of first runner up (second highest likelihood). **b)** Percentage of original classifications in runners up for incorrectly classified cells. **c)** Time taken in seconds to classify all cells in dataset for the different classifier. **d-e)**. Bootstrapped runs of classification of cell types with 5-30% of marker genes with inputed noise **(d)** or downsampled **(e)**. Micro F1-score shown.

Another feature of BLC is that it also provides the second and third most likely cell type, so called “runners up”. In fact, for cells wrongly annotated by BLC the original annotation was first runner up in most cases (84.3%), sometimes a second runner up (10.5%), but on a few occasions it was not included at all (5.2%) (**Figure 5b**). This suggests that when BLC returns a lower certainty level for the primary annotation, one can inspect the runner up cells more closely. Again, compared to scPreds “runners up”, BLC showed a greater inclusion of original cell classifications among the runners up (**Figure 5b**).

Another highly important aspect of BLC is that even though it is performed on a cell-by-cell basis, the time spent on cell annotation is less than for the clustering algorithm and considerably less than for scPred (**Figure 5c**). BLC was faster than scType and scPred for all 4 datasets tested. Compared to scPred, it was approximately 5.7 times as fast for the largest dataset from Oelen et al, consisting of 39,172 cells (104 seconds compared to 592 seconds), and about 3.5 times faster for the smallest dataset (SmartSeq data, 2841 cells). The timing was started immediately prior to initial normalization of the data, to provide a complete overview of how long both scripts took to annotate cells.

As well as taking less time, BLC also utilized less data resources for classifying cells compared to scType and scPred (**Extended Figure 4b**). Similarly, concerning the training of BLC and scPred models, BLC was significantly more efficient than scPred, completing training in just 33 minutes versus 6.5 hours for scPred, with mean memory allocations of 21 GB for BLC and 33 GB for scPred, respectively (**Extended Figure 4c**). This efficiency is increasingly critical as the number of cells and the depth of sequencing continues to grow with advancing scRNA-seq technology.

Finally, we wanted to challenge BLC using noisy and down-sampled data. For the noisy data, we conducted 100 iterations of data classification, each time introducing noise to 5-30% of the marker genes using a Poisson distribution with a lambda (λ) value of 1. In each iteration, a random set of marker genes was subjected to noise, thereby ensuring a robust assessment of the resistance of BLC to noise. BLC maintained a stable performance until 20% of the marker genes were influenced by noise **(Figure 5d, Extended Figure 5)**.

We followed a similar schedule for the down-sampling of genes, with subsampling and deletion of 5-30% of the marker genes that the classifier had chosen. Again, BLC was robust against a down-sampling of genes of up to 20% (**Figure 5e, Extended Figure 5**), which is particularly useful for classifying scRNA-seq data sequenced with less read depth, enabling accurate cell type identification even when marker gene expression is somewhat limited.

In summary, BLC provides reliable annotations of cell classifications with good assessment of when to doubt itself, often with well-founded runners up. Additionally, it is faster and utilizes considerably less data resources than other models showing similar classification accuracy, an aspect of great importance considering the increase in scRNA-seq dataset sizes. Furthermore, BLC shows resistance towards noisy genes and down-sampled data.

## Discussion

This study introduces the BayeLeafClassifier (BLC), a novel tool for finding marker genes that can separate cell types or cell substates for scRNA-seq data analysis, and further train a model to classify previously unseen data into training categories. BLC provides highly accurate and robust results in both perturbed and unperturbed cells and offers a probabilistic way to consider cell annotations. This probability can be further examined, providing an additional layer of information for interpretation Further, the method is resistant to *in silico* data perturbations, is faster than traditional clustering methods and requires less computational resources.

A notable strength of BLC is its sensitivity towards less common cell types and closely related cell types, which traditional clustering methods often struggle to accurately differentiate. BLC effectively discerns such populations, as evidenced by its superior performance compared to scType, especially in distinguishing cell types such as CD4^+^ T cells and CD8^+^ T cells, NK cells, and various dendritic cells. We observed that the majority of incorrectly annotated cells were associated with lower uncertainty levels, contrasting with higher certainty levels for correct annotations. This feature of BLC not only aids identification of potential errors, but also indicates a sophisticated self-assessment capability within the model, a valuable characteristic enabling users to identify and scrutinize potentially ambiguous cases.

The efficiency of BLC is further underscored by its use of approximately 140 marker genes to separate different cell types, a significantly smaller number than the 2000-3000 typically employed in defining transcriptional signatures in clustering algorithms. Despite the reduced number of markers, BLC based on EMD successfully identifies genes that preserve enough information about a cell type, demonstrating that a concise set of markers can be as effective as a more extensive set, if judiciously selected. This allows for an efficient classification and training of the model, both regarding time spent and computational resources, particularly evident when benchmarked against a cell-by-cell classifier working in a reduced dimensional space.

One important consideration concerning the performance of scPred is that it was trained on approximately 300,000 cells instead of the 400,000 cells used for BLC training. While increasing the size of the dataset might improve F1-scores, this was not possible due to computational limitations. With more computational resources it might be possible to train the scPred model on a larger dataset, and thereby perhaps reach better F1-scores to outperform BLC. However, in such a scenario, BLC would achieve an even greater advantage in terms of time spent for model training and computational resources required, and certainty-levels would still be more reliable with BLC.

Using non-linear quantile transformation instead of the more commonly used Z-score transformation is associated with several advantages in the creation of BLC. First, the EMD becomes comparable between different genes since all are expected to be uniformly distributed between 0 and 1. This ensures that each gene contributes equally to the EMD and allows a more balanced comparison between different marker genes. Secondly, it becomes less sensitive to outliers compared to Z-score transformation. This provides an advantage when classifying previously unseen datasets where differences in read depths could skew the distribution of a marker gene, and where quantile transformation could handle this skew more effectively. However, it could also skew the unseen data if the training and test datasets originate from different populations (e.g. training on a full PBMC dataset, but classifying using only lymphocytes). It is therefore important that the expected populations in the test dataset are the same as for the training dataset.

While BLC marks a significant step forward, it is not without limitations. The fact that the model relies on existing annotations for training could introduce biases or errors inherent to the primary data. An interesting caveat is that the original cell type classifications that BLC was trained on were classified using clustering algorithms. It could therefore be a justified criticism that the classifier is suffering from the same issues as clustering algorithms since it was trained to recognize these cluster-defined cell types. However, here we show that BLC identifies cells that according to the original annotation and the benchmarked clustering tool were wrongly labeled, but where canonical marker genes were strongly in favor of the BLC annotation. Therefore, it seems that BLC does not suffer from the oversimplification of transcriptional signatures as clustering does, nor the distortion of the data.

Additionally, the performance of the classifier, like any computational tool, is dependent on the quality and representative nature of the input data. Future iterations of BLC could focus on enhancing its ability to learn from a wider range of data types, including more challenging and even less well-characterized cell states or cell type subsets. This can readily be performed as BLC is easily modified to find other groupings of interest in scRNA-seq. As BLC is a general framework to find both the best marker genes and to classify previously unseen datasets, it is very versatile and can be used in many different contexts.

As cells develop in a hierarchical fashion, closely related cell types will have very similar transcriptional profiles with specific marker genes separating them, as is the case for CD4^+^ T cells and CD8^+^ T cells. One could make the case that a more holistic view of the whole transcriptome is important for cell types that are further away from each other in the developmental trajectory, such as dendritic cells and CD8^+^ T cells. However, we do not see any major mislabeling of these cells using our BLC classifier, even though we only examine ∼140 marker genes for each cell pair, and do not firstly cluster the cells on e.g. 2000 highly variable genes. This suggests that the marker genes that we found are robust for any inference between different cell types, no matter how close or distant they are on the developmental trajectory.

In conclusion, BLC represents a powerful and reliable tool for cell type classification of single-cell RNA sequencing data. Its ability to deliver accurate, stable and sensitive classifications, coupled with its efficient use of marker genes, self-assessment of prediction probability and enabling further inspection of the certainty in its annotations, make it a valuable asset in the toolkit of computational biologists and researchers in the field of genomics.

## Data availability

No new data was generated for this publication, all datasets are public and have been published in other papers. See supplementary material and methods for data collection.

## Code availability

BayeLeafClassifier is available on github at https://github.com/rikfor/BayeLeafClassifier with all code used to generate the figures in this article and notebooks for how to train new classifiers and predicting using newly trained classifiers. The classifier used for this article is available at https://drive.google.com/drive/u/0/folders/1--eGmL9U-LCvs_G_EdhPEQDKIjZbvWuI.

## Acknowledgements

The authors thank members of the Brodin lab and Dimarogonas laboratories for helpful discussions and input on the work. This work was supported by a grant from the Knut and Alice Wallenberg Foundation to SciLifeLab for research in Data-driven Life Science, DDLS (KAW 2020.0239) program. The Brodin laboratory is also supported by HORIZON HLTH-2021-DISEASE-04 program under grant agreement 01057100 (UNDINE), the Swedish research council (2019-01495, 2020-06190, 2020-02889, 2021-06529, 2021-05450, 2022-01567) and Knut & Alice Wallenberg Foundation (VC-2021-0026). Göran Gustafsson Foundation (GG20200040). Swedish Society for Medical Research (SSMF, CG-22-0148-H-02).

