## Supplementary material for "Cluster-free annotation of single cells using Earth mover’s distance-based classification": Methods and extended figures

**Construction of EMD-distribution calculator**

EMD marker genes were found using our *BLC_classification_lib.py*, which can be found on github at https://github.com/rikfor/BayeLeafClassifier.

The EMD measures the work needed to transform one probability distribution into another and that the work involved is directly proportional to the amount of earth moved.

For 1-dimensional probability distribution functions, the calculation of earth mover's distance is as follows:

$$D^{g}_{EMD}\left(CT(A), CT(B)\right) = \int_{-\infty}^{\infty} \left| \int_{-\infty}^{v} \left( P\left(x_g = v' \middle| CT(A)\right) - P\left(x_g = v' \middle| CT(B)\right)\right) dv' \right| dv$$

where P(xg | CT(A)) and P(xg | CT(B)) are respectively the probability distributions of the g-th gene in cells from the classes CT A and CT B. We adopt a binning approach where we divide each continuous probability distribution into n=1000 bins to obtain a discrete approximation.

In other words, genes distributions from two cell types are combined into one pandas DataFrame which undergoes a cumulative summation, which represents the aggregated differences in gene expression up to each point in the distribution. The cumulative function is then normalized by the "carry length", here defined as the difference in the index values between successive rows in the merged pandas DataFrame and the "total_length", which is the total length of the DataFrame. This results in a final value representing the distance between the gene distributions of two cell types.

For each gene and each cell type pair, EMD is calculated and the genes with the highest distance over a user-set threshold (0.4 in this paper) were selected, with a maximum number of genes to look at included (selecting the highest ones), and a minimum genes selection that, if necessary, goes below the threshold. Which cell type the gene provide a marker for was selected by comparing median expression. For visualization, see **Extended Figure 6**.

**Integrating Bayesian classifier with Random Forest**

For BLC, a naïve Bayesian classifier was used as a root node for each decision tree in the Random Forest. This employs Bayes' theorem to compute the posterior probability of a cell being a certain cell type, which is given by:

$$P(C_k | x_1, x_2, \dots, x_n) = \frac{P(C_k) \prod_{i=1}^{n} P(x_i | C_k)}{P(x_1, x_2, \dots, x_n)}$$

where $P(C_k | x_1, x_2, \dots, x_n)$ is the posterior probability of class Ck given the features x1,x2, ..., xn. P(Ck): The prior probability of class Ck, which is the general likelihood of class Ck occurring in the dataset. $\prod_{i=1}^{n} P(x_i | C_k)$ is the likelihood of observing the set of features x1,x2,...,xn given class Ck. This is calculated as the product of the probabilities of each feature xi given class Ck, assuming each feature contributes independently (hence the naivety). P(x1 ,x2,...,xn) is the probability of observing the set of features x1,x2,...,xn regardless of the class. This acts as a normalizing constant to ensure that the posterior probabilities across all classes sum up to 1.

Following the Bayesian classification at the root, decision trees in the Random Forest are grown to their maximum extent without pruning, with splits based on Gini impurity.

**BayeLeafClassifier cell type annotation and certainty**

During the cell-by-cell annotation process, we compare each cell types two by two, and count the number of estimators that are in favor of one cell type over the other, such $Y_{RF} = \mathrm{mode}\{Y_1, Y_2, \ldots, Y_N\}$, Y being a decision tree. This can be likened to a voting system where each of these estimators casts a 'vote' for one of the two cell types. The final annotation for a given cell is determined by majority rule; that is, the cell type that receives the highest number of votes from the estimators is selected as the label for that cell. After going through all the cell pairs, we again take the majority-winner between all of the cell pair comparison, ending in a majority decision with the labeling of a cell.

Furthermore, we quantified the certainty of that annotation by examining the proportion of votes for each cell type. For instance, if we have a total of 500 estimators and 400 of them votes for cell type X, and 100 for cell type Y, then it is a 400/500 = 80% certainty that it is cell type X. This certainty is reflected for the second and third most likely cell type, named "likelihood_vs_second_ct" and "likelihood_vs_third_ct" in the prediction output. For "overall_likelihood", we take the mean of three smallest likelihood, as an estimation of the broader perspective for the cell type assigned.

**Construction of training dataset for BayeLeafClassifier**

In order to train BayeLeafClassifier on representative data of PBMCs, we collected 422,587 cells from 12 different studies of both stimulated and unstimulated cells: 2 of them from Chan-Zuckerberg Initiative CELLxGENE; *A blood atlas of COVID-19 defines hallmarks of disease severity and specificity* (1) and *Systems biological assessment of immunity to mild versus severe COVID-19 infection in humans (2)*.

5 of the datasets were downloaded from Gene Expression Ominbus (GEO): Single-cell RNA-seq reveals cell type–specific molecular and genetic associations to lupus (CLUES) (GSE174188) (3) , Time-resolved Systems Immunology Reveals a Late Juncture Linked to Fatal COVID-19 (GSE161918) (4) , Longitudinal single-cell epitope and RNA-sequencing reveals the immunological impact of type 1 interferon autoantibodies in critical COVID-19 (GSE168453) (5), COVID-19 immune features revealed by a large-scale single-cell transcriptome atlas (GSE158055) (6) , Single-cell Transcriptomic Landscape of Human Blood Cells (GSE149938) (7).

*RNA-seq of RBC depleted whole blood from COVID-19 patients and controls* (8) was collected from Biostudies (https://www.ebi.ac.uk/biostudies/).

*Local and systemic responses to SARS-CoV-2 infection in children and adults* (9) and *Single-cell multi-omics analysis of the immune response in COVID-19* (10) were collected from https://www.covid19cellatlas.org/. Zheng et al dataset (11) was collected from 10X genomics datasets portal https://www.10xgenomics.com/datasets (as "Fresh 68k PBMCs (Donor A)).

Finally, *Single cell immune profiling reveals distinct immune response in asymptomatic COVID-19 patients* (12) was downloaded from CNGB Nucleotide Sequence Archive (CNSA), accession number CNP0001250.

The studies were of different sizes, so in order not bias BayeLeafClassifier toward any dataset we downsampled the larger ones by randomly selecting cells, however we did not downsample rarer cell types such plasmacytoid dendritic cells, dendritic cells and plasma B cells, which were retained in full to ensure a rich dataset for these rare cell types.

**BLC workflow options**

The naïve Bayesian Classifier implemented through *naive_bayes* implementation from sklearn (13) and was trained with default parameters, however default values for variance smoothing can be changed according to users preference (default = 1e-9). For Random Forest, again multiple default values can be changed according to users' preferences, but the following set of parameters was changed in the training of BLC: The maximum depth of the trees in the Random Forest was set at 40, a choice made to balance the complexity of the model against the risk of overfitting. To ensure robustness in the model, 200 estimators were used, defining the number of trees in the forest. We also set the minimum number of samples required to be at a leaf node to 40, ensuring sufficient data points for each decision path in the trees, and a random_state = 0.

Additionally, BLC allows the user to control for how many features from EMD-calculation to be input into model training. This includes balanced_ct_genes, which is an option if the user want to enforce an equal balance in the number of marker genes from both cell types. In this article, balanced_ct_genes was set to False. BLC also allows to change the maximum and minimum number of genes to be considered. Here, maximum number of genes for each cell type were set at 70 and minimum genes to consider was set to 5, providing a range that captures sufficient genomic information while avoiding excessive computational complexity. Finally, a threshold of EMD >= 0.4 was chosen for the earth mover's distance (EMD), ensuring a precise and meaningful differentiation between cell types based on their gene expression profiles, a setting that is also easy to change for the user.

The full set of parameters options can further be found in the extensive documentation of sklearn for RandomForestClassifier, GaussianNB and at the user's tutorial found in github associated with this paper.

**Training of scPred**

scPred was trained on 321,000 cells, downsampled from the original data of 422,587 cells. Due to computational limitations and that scPred requires substantial more amount of computational power than BLC, we could not train scPred on more than the ~300,000 cells. scPred was trained according to their official tutorial, with allowance for parallelization when training to speed up the training. The script for training scPred is available at github repository linked with this manuscript.

**Systematic comparison in publicly available dataset**

In benchmarking against scType, the cells were first clustered through the Seurat workflow (version 5.0.1). Default parameters was used, but varied in the number of dimensions in *Seurat::FindNeighbors* and *Seurat::RunUMAP* (14) based on the guidance from ElbowPlot. After finding the appropriate dimensions, we executed the entire scType annotation script again to accurately measure the time required for cell annotation. The script used for scType annotations is available in the GitHub repository associated with this paper. Given that

scType provides annotations at a finer resolution than our classifier was trained for (e.g., distinguishing between "CD4+ memory T cells" and "CD4+ naïve T cells", instead of labeling both as "CD4+ T cells"), we adapted the scType annotations to match our broader classification categories. For instance, annotations like "CD4+ memory T cell" were relabeled to a more general "CD4T" category. Additionally, cells annotated as "Macrophages" by scType, which more accurately correspondeds to "Non-classical Monocyte" or "Classical Monocyte" were relabeled to their original annotations, as the "Macrophage"-annotation per se is not entirely incorrect. Similar relabeling to cells' original annotations was done for cells marked by scType as Interferon Stimulated Genes (ISG) + cells, or Unknown cell type. As BLC does not have Unknown category by default this would be an unfair comparison.
The complete details of this reannotation process can be found in scripts for *Figure 2* on provided github.

**Downsampling and noise imputation of genes**
To evaluate the robustness of our classifier against both technical noise and data downsampling, we conducted a series of systematic tests. For assessing noise resistance, we introduced randomness into the marker gene values, by adding values drawn from a Poisson distribution with $\lambda = 1$. The noise was added to an incrementally increased proportion of the marker genes, ranging from 5% to 30%, thus we evaluated the classifier's performance under gradually escalating noise levels in marker genes.

In parallel, we tested the classifiers resistance to downsampling, which simulates scenarios of data loss or incomplete marker gene information. Following a similar schedule as for the noise-resistance test, we systematically removed 5% to 30% of the marker genes by setting them equal to zero.

Both scripts for testing noise imputation and downsampling of genes resistance are available on github at https://github.com/rikfor/BayeLeafClassifier.


1. Ahern DJ, Ai Z, Ainsworth M, Allan C, Allcock A, Angus B, et al. A blood atlas of COVID-19 defines hallmarks of disease severity and specificity. Cell. 2022 Mar;185(5):916-938.e58.

2. Arunachalam PS, Wimmers F, Mok CKP, Perera RAPM, Scott M, Hagan T, et al. Systems biological assessment of immunity to mild versus severe COVID-19 infection in humans. Science. 2020 Sep 4;369(6508):1210–20.

3. Perez RK, Gordon MG, Subramaniam M, Kim MC, Hartoularos GC, Targ S, et al. Single-cell RNA-seq reveals cell type–specific molecular and genetic associations to lupus. Science. 2022 Apr 8;376(6589):eabf1970.

4. Liu C, Martins AJ, Lau WW, Rachmaninoff N, Chen J, Imberti L, et al. Time-resolved systems immunology reveals a late juncture linked to fatal COVID-19. Cell. 2021 Apr;184(7):1836-1857.e22.

5. Van Der Wijst MGP, Vazquez SE, Hartoularos GC, Bastard P, Grant T, Bueno R, et al. Longitudinal single-cell epitope and RNA-sequencing reveals the immunological impact

of type 1 interferon autoantibodies in critical COVID-19 [Internet]. Immunology; 2021 Mar [cited 2024 Feb 11]. Available from: http://biorxiv.org/lookup/doi/10.1101/2021.03.09.434529

13. Pedregosa F, Varoquaux G, Gramfort A, Michel V, Thirion B, Grisel O, et al. Scikit-learn: Machine Learning in Python. MACHINE LEARNING IN PYTHON.

14. Butler A, Hoffman P, Smibert P, Papalexi E, Satija R. Integrating single-cell transcriptomic data across different conditions, technologies, and species. Nat Biotechnol. 2018 May;36(5):411–20.

### *Extended Figures*

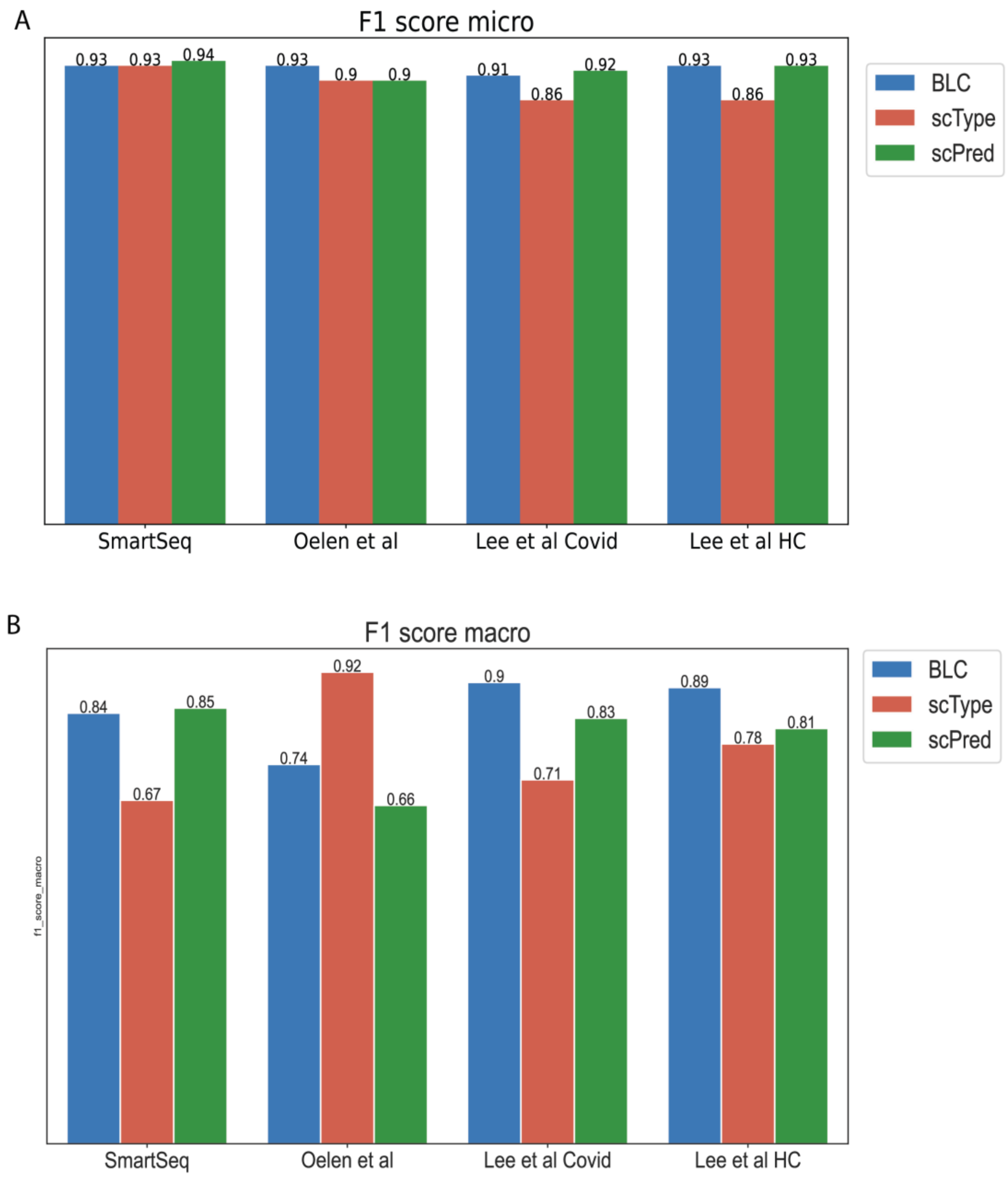


Extended Figure 1. A) F1-micro score for all classified dataset. B) F1-macro score for all classified datasets.

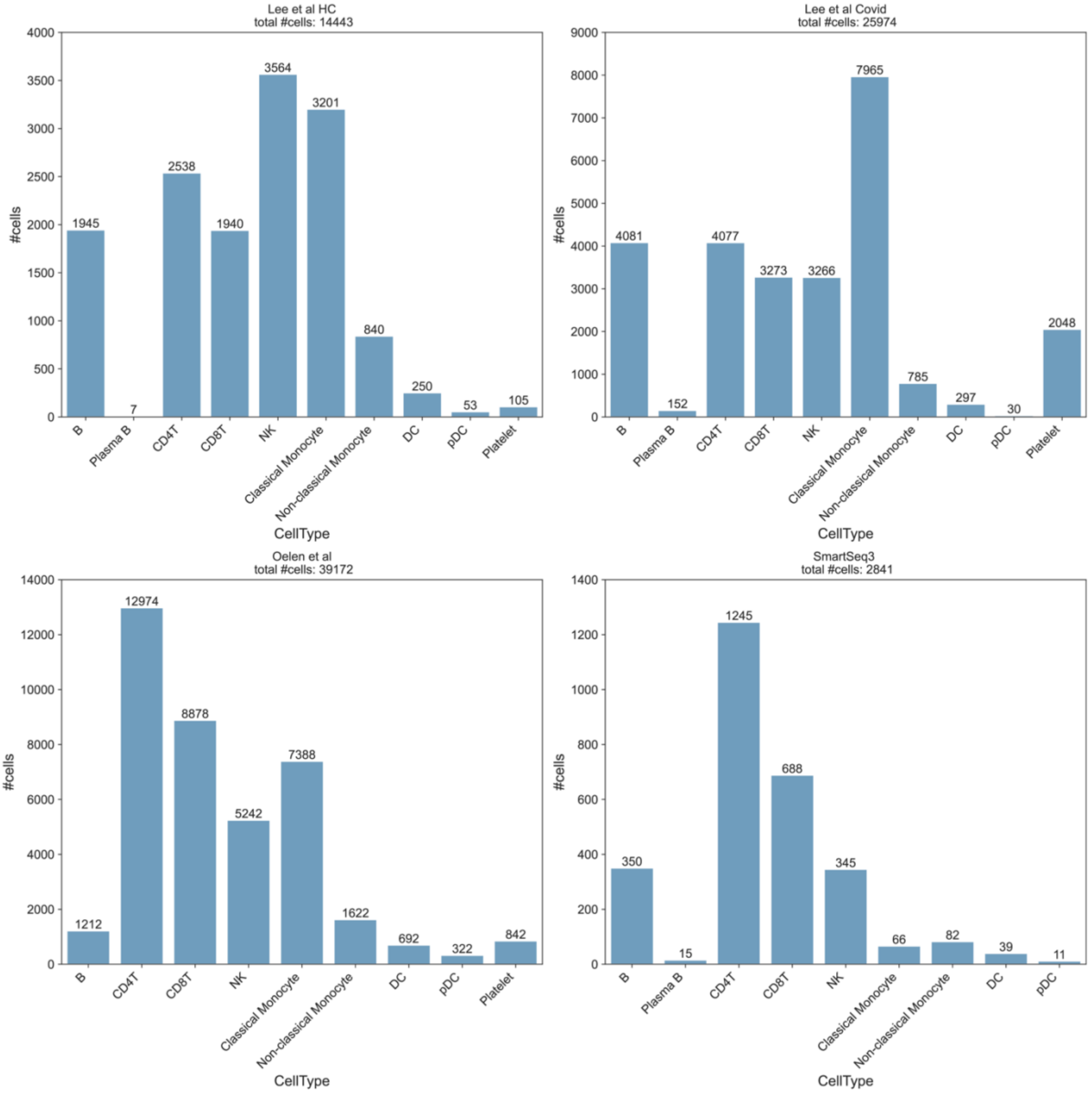


Extended Figure 2. Number of cells for each cell type in every dataset tested.

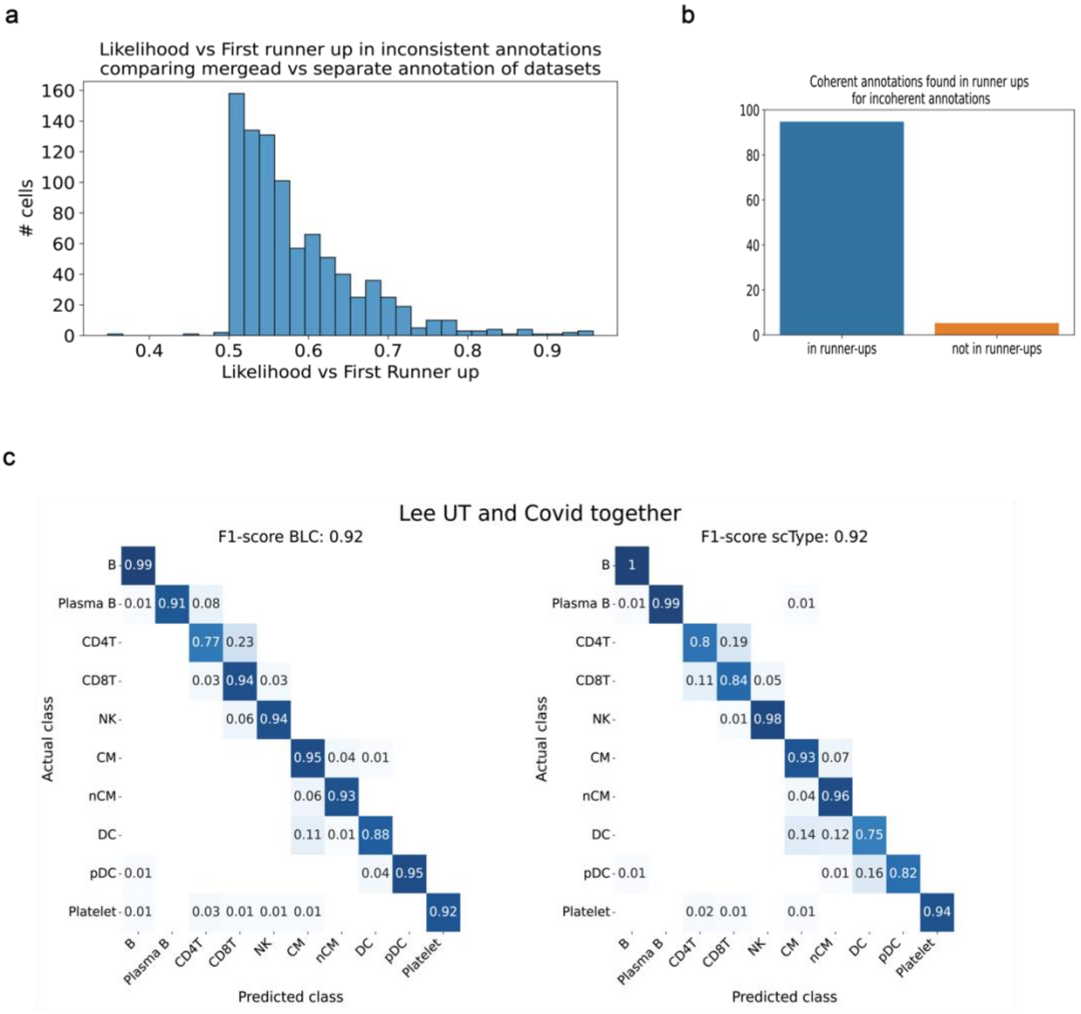


Extended Figure 3. **(a)** Likelihood between selected annotation and first runner up for inconsistent annotations when classifying Lee et al data separate and together. **(b)** Percentage of consistent annotations found in runner ups for inconsistent annotations. **(c)** Heatmaps showing micro F1-score and percentage correct annotations of each cell type in dataset, when classifying the two datasets (Covid-19 and helathy controls (HC)) from Lee et al together. BLC left panel, scType right panel.

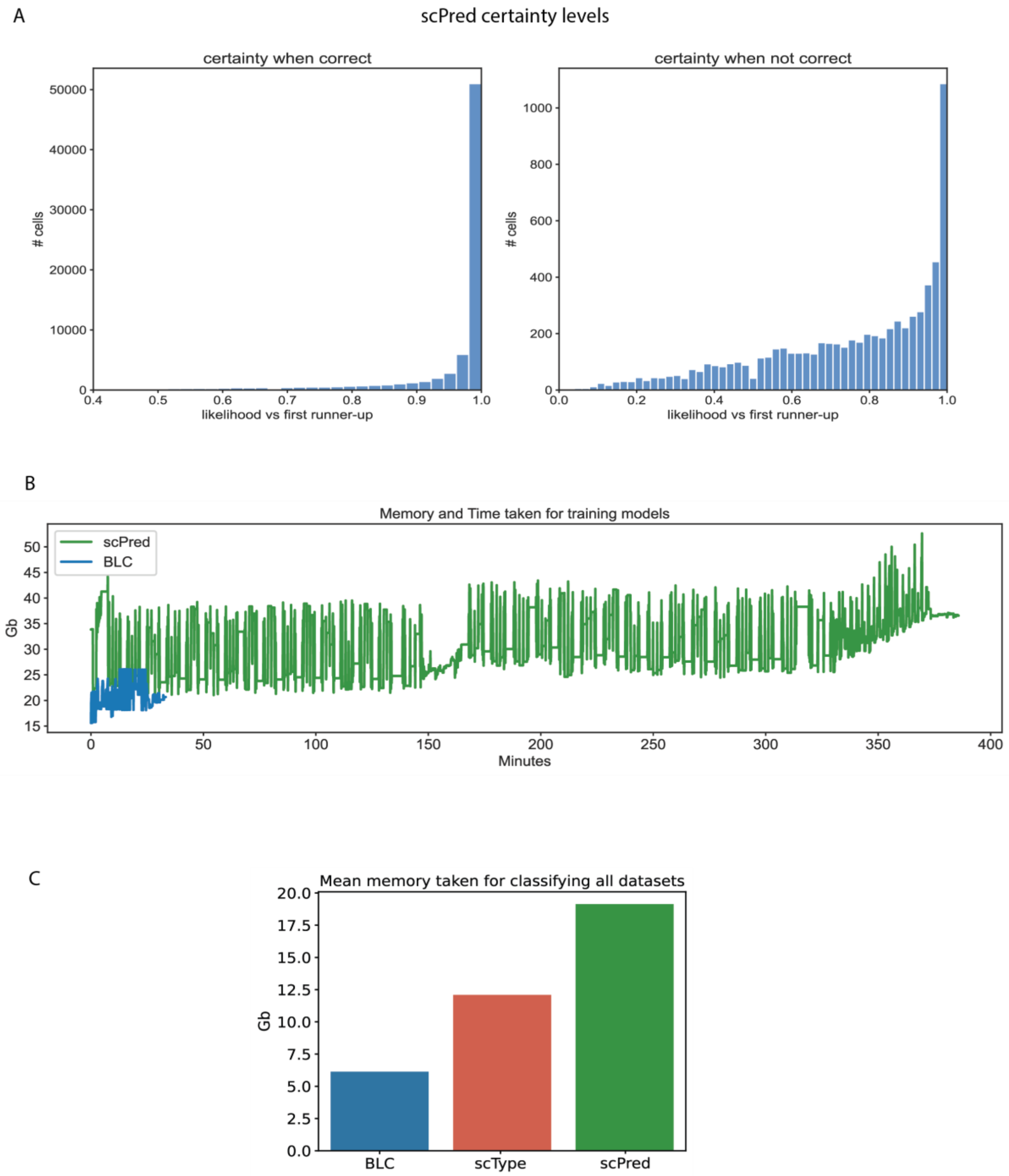


**Extended Figure 4.** A) scPreds certainty level for when correct (left panel) and when not corect (right panel). B) Comparison of memory used in gigabytes (Gb) (y-axis) and time taken (x-axis) during training of scPred and BLC. C) Mean memory taken in gigabytes (Gb) for when classifying all datasets.

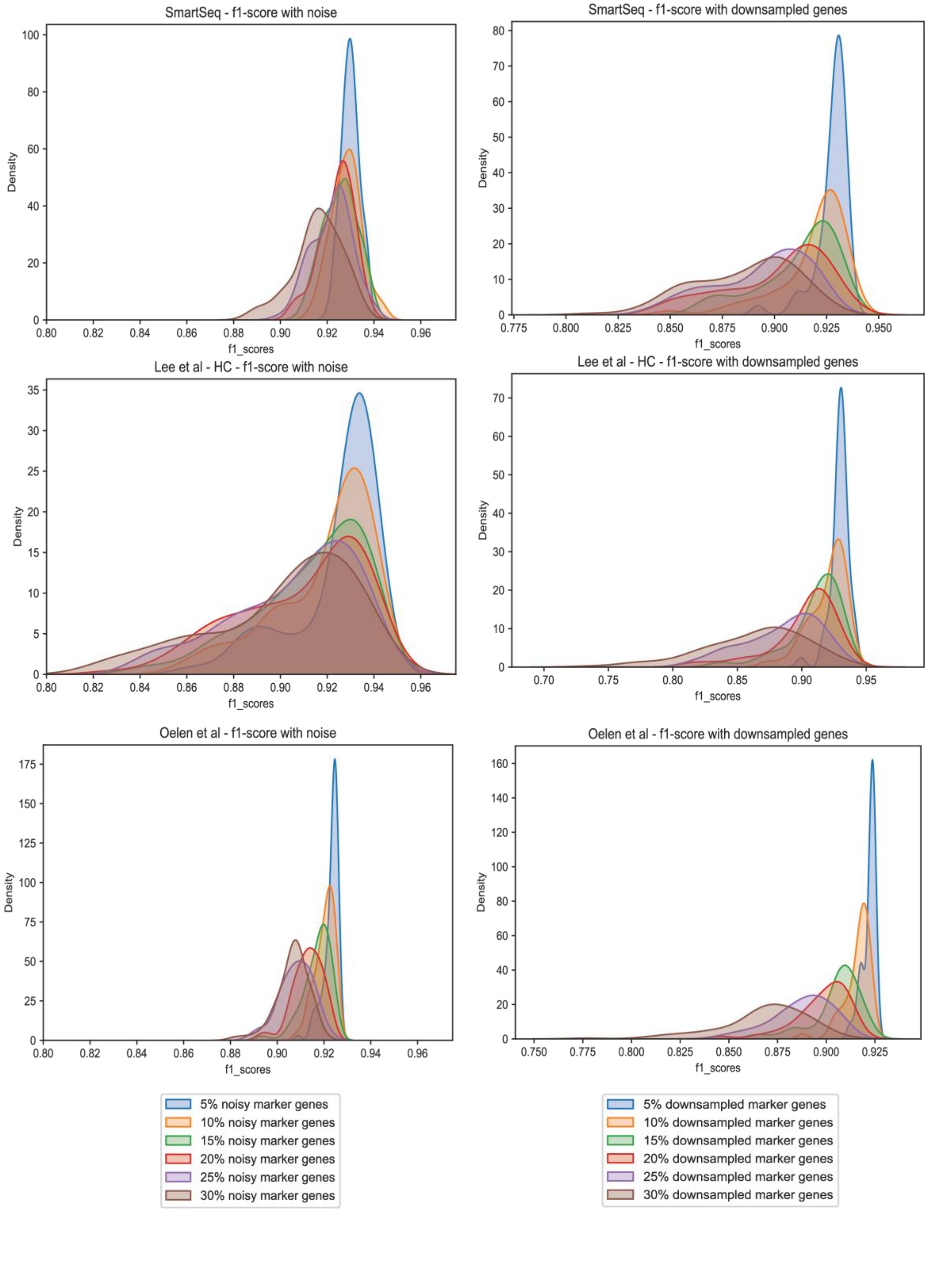


Extended Figure 5. Bootstrapped runs of classification of cell types with 5-30 % of marker genes with inputed noise (left column) or downsampled (right column). Micro F1-score shown.

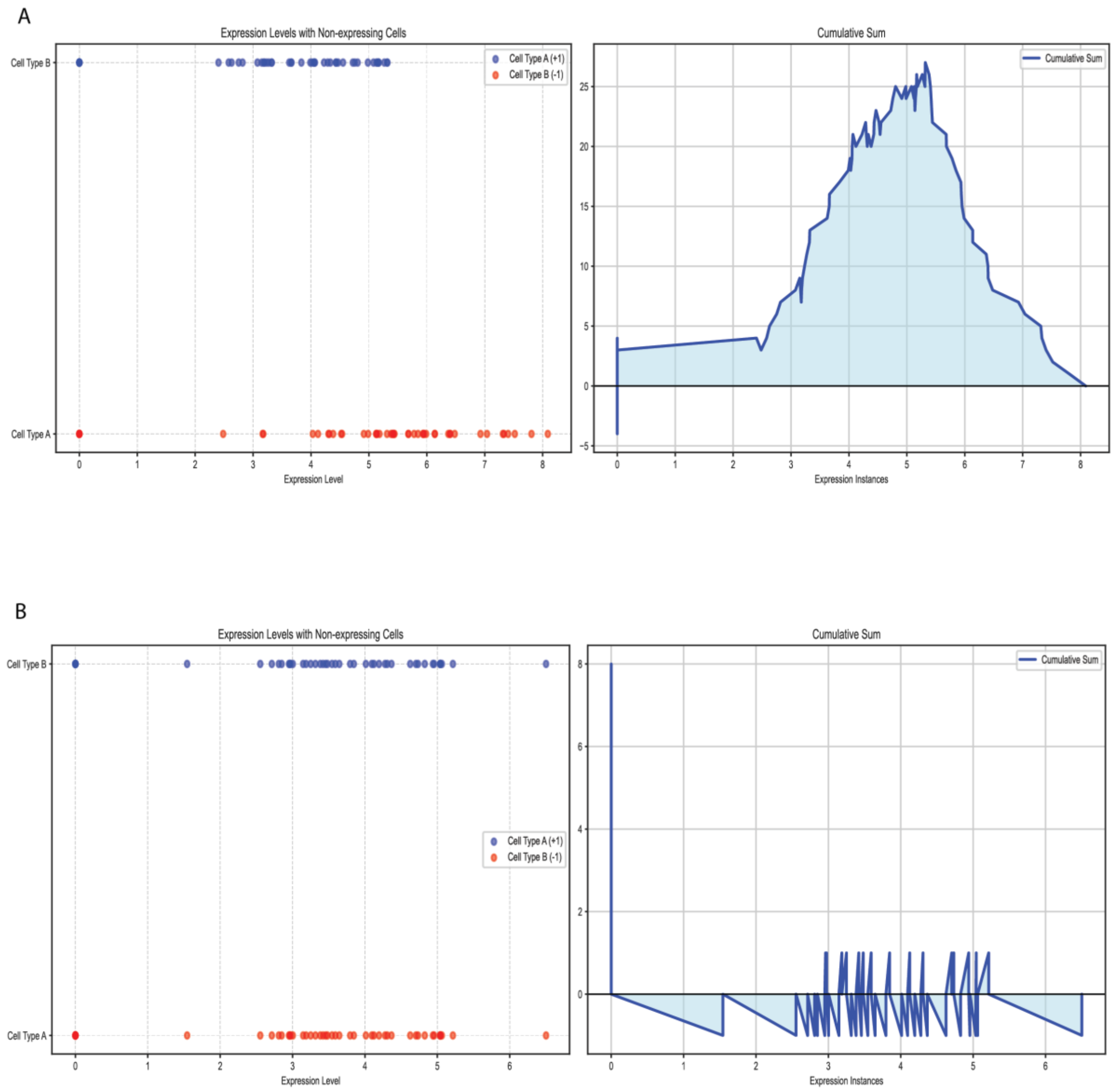


Extended Figure 6:
Given the same number of cells from cell type 1 and cell type 2, we measure the EMD for a gene through an indexed series where the index is the level of expression by each cell and the value is +1 if this is an expression from a type A cell and -1 if it is an expression from type B cell. The total sum is as a result zero. The cumulative sum can be positive or negative at each expression value depending on the difference of expression up to that point. The EMD between the two expression distributions is the Integration of the absolute value of the cumulative sum. A) showing high EMD B) showing low EMD
